# Inverse-scattering in biological samples via beam-propagation

**DOI:** 10.1101/2025.08.17.670744

**Authors:** Jeongsoo Kim, Blythe Bolton, Khashayar Moshksayan, Rishika Khanna, Mary E. Swartz, Michał Ziemczonok, Mohini Kamra, Karin A. Jorn, Sapun H. Parekh, Małgorzata Kujawińska, Johann Eberhart, Elif Sarinay Cenik, Adela Ben-Yakar, Shwetadwip Chowdhury

## Abstract

Multiple scattering limits optical imaging in thick biological samples by scrambling sample-specific information. Physics-based inverse-scattering methods aim to computationally recover this information, often using non-convex optimization to reconstruct the scatter-corrected sample. However, this non-convexity can lead to inaccurate reconstructions, especially in highly scattering samples. Here, we show that various implementation strategies for even the same inverse-scattering method significantly affect reconstruction quality. We demonstrate this using multi-slice beam propagation (MSBP), a relatively simple nonconvex inverse-scattering method that reconstructs a scattering sample’s 3D refractive-index (RI). By systematically conducting MSBP-based inverse-scattering on both phantoms and biological samples, we showed that an amplitude-only cost function in the inverse-solver, combined with angular and defocus diversity in the scattering measurements, enabled high-quality, fully-volumetric RI imaging. This approach achieved subcellular resolution and label-free 3D contrast across diverse, multiple-scattering samples. These results lay the groundwork for robust use of inverse-scattering techniques to achieve biologically interpretable 3D imaging in increasingly thick, multicellular samples, introducing a new paradigm for deep-tissue computational imaging.

## Introduction

Optical microscopy is a major research tool in the biosciences and enables both morphological and molecular imaging at subcellular spatial resolutions and high speeds. However, a major challenge is optical scattering, which scrambles sample-specific information and degrades imaging quality starting at depths of even just a few tens of microns into biological tissue. Thus, classical microscope techniques are typically used to image only outer superficial tissue layers, or specific components isolated from within tissue and prepared *in-vitro*.

In response to this challenge, deep-imaging techniques (e.g., multiphoton microscopy, optical coherence tomography, etc) have emerged that extend imaging depths to the order of multiple hundreds of microns, made possible by selectively exciting or detecting with *only* unscattered (i.e., ballistic) light. These deep-tissue imaging techniques have found enormous success in the biosciences in enabling up to millimeter-scale imaging depths in fields such as neuroscience^1-3^, developmental biology^4-6^ and biomedicine^7-9^. However, the ballistic component of the total signal decreases exponentially with depth – as a result, these deep-imaging techniques are still fundamentally constrained by scattering.

Interestingly, though scattered light appears random, it deterministically encodes sample-specific information. Furthermore, light can scatter through several millimeters in tissue before being absorbed. Thus, if one could *fully* unscramble the effects of scattering, one could in theory gain access to an imaging volume with more than an order of magnitude longer imaging depth than that of current state-of-the-art deep-imaging techniques. Though this has yet to be achieved, it remains a powerful motivation that drives significant excitement and ongoing efforts to develop methods for correcting optical scattering.

In the biosciences, scatter correction is often exemplified by adaptive optics (AO). AO typically begins by sensing wavefront distortions from a point-emitting source (i.e., a guidestar) within tissue, followed by applying physical wavefront-shaping elements to correct those distortions. This enhances image quality in the surrounding region, whose lateral extent (i.e., isoplanatic patch) is determined by intrinsic tissue correlations. AO has seen broad adoption across various biological imaging fields, such as brain science^10-12^, vision science^13-15^ and cell biology^16-18^, but faces key limitations: (1) it requires time-intensive calibration localized to small tissue regions, hindering scalability to large 3D volumes; (2) guidestar placement within tissue is technically challenging. While “sensorless” AO methods avoid this, they are slower and risk photobleaching^19,20^; (3) AO primarily addresses low-order scattering, while deeper tissue imaging suffers from high-order, multiple-scattering effects. Though newer wavefront-shaping methods can correct for these, they often rely on complex setups and remain less widely adopted in biosciences^21-23^.

A separate class of optical scatter-correction techniques has emerged that leverages computational methods to reconstruct fully volumetric, scatter-corrected 3D images. These frameworks typically use physics-based scattering models to describe optical paths that light takes within scattering media. Importantly, these models need to be invertible so that a 3D image of the sample can be reconstructed from an input set of raw scattering measurements via either analytic inversion or iterative optimization. This process is broadly referred to as solving the inverse-scattering problem and holds special importance even in fields outside optical imaging^24-27^. Compared to AO, which aims to characterize the *accumulated* scattering effects from a limited region within the sample, inverse-scattering methods seek to characterize the sample’s intrinsic scattering properties *fully in 3D*. This is popularly accomplished by mapping out the sample’s 3D refractive-index (RI), which fully characterizes the sample’s scattering behavior due to its direct mathematical equivalence to scattering potential. Notably, RI is also a powerful source of endogenous morphological contrast – thus, mapping out a sample’s 3D RI naturally provides scatter-corrected, label-free, and quantitative tomography of the sample’s 3D morphology. Because this method does not require guidestars, it is particularly promising for non-invasive imaging applications.

Various inverse-scattering methods have been developed to reconstruct a sample’s 3D RI from measured scattering data. In the simplest case when a sample is *weakly* scattering, scattering measurements can be *linearly* related to the sample’s 3D RI. This enables 3D RI to be directly reconstructed from measurement data via simple linear inversion. This strategy found early and significant success in imaging applications involving *thin* biological samples (e.g., mono-layer cell distributions, small multicellular samples, etc) where imaging depths were typically a few tens of microns. Multiple groups have introduced variations of this strategy to reconstruct 3D RI under a variety of imaging configurations, such as interferometric^28,29^ versus non-interferometric detection^30,31^, monochromatic^32,33^ versus broadband illumination^34,35^, illumination-angle^36,37^ versus axial-defocus scanning^38,39^, planar^40,41^ versus patterned illumination^42,43^, etc. Many of these methods have matured to the point where they are robust enough for studying sensitive biological processes, which often demand long-term stable imaging with minimal phototoxicity. Recent commercialization efforts (e.g., Tomocube, Nanolive, PhiOptics) have further accelerated the global adoption of RI tomography, paving the way for its widespread use in research labs worldwide.

However, a major limitation of the inverse-scattering methods described above is their reliance on the assumption that the sample is *weak* scattering. This assumption linearizes the inverse-scattering problem enabling easy reconstruction of *thin* samples, such as monolayer cell samples. For thicker and larger-scale multicellular samples, which contain significant *multiple* scattering, recent studies have introduced partially-coherent RI tomography techniques that employ coherence gating to selectively detect only the weak-scattered component of the total scattered light^43-45^. This enables the continued use of weak-scattering assumptions for 3D RI reconstruction in multicellular samples. However, as sample thickness increases, the weakly scattered signal may be rapidly overwhelmed by multiple scattering, eventually rendering it nearly undetectable.

More sophisticated models have been introduced that mathematically account for some degree of multiple scattering. For example, beam propagation methods – originally developed decades ago for modeling light transmission through atmospheric turbulence^46^, graded-index fibers^47^, microlenses^48^, etc – have recently been adapted into optical microscopy for 3D inverse-scattering and RI reconstruction of multicellular organisms such as *C. elegans* worms^49,50^, mouse embryos^51,52^, or zebrafish embryos^53^, where optical thicknesses can range to >60 microns. Independent work by Osnabrugge *et al*.^54^ introduced a separate inverse-scattering framework based on a modified Born series, which was later demonstrated for bio-imaging by Lee *et al*.^55^ and Li *et al*.^56^, who show RI reconstructions in tissue sections, microalgae, and *C. elegans* worms. A series of works by Yasuhiko *et al*.^57-59^ introduced another separate inverse-scattering strategy that computationally suppressed scatter effects via either coherent or incoherent accumulation. These synthetically generated scatter-suppressed measurements could then be related to the sample’s 3D RI distribution via deconvolution. Biological demonstrations were conducted with organoid samples, where imaging results included multicellular human hepatoma organoids and human brain glioblastoma spheroids with thicknesses greater than 130 microns.

Notably, despite promising demonstrations, these types of inverse multiple-scattering methods have yet to gain widespread adoption in the basic-science community. A major reason is likely that their performance has not yet been characterized across diverse, real-world biological samples that exhibit varying degrees of multiple-scattering. Moreover, the fundamental limits of these methods remain largely unexplored. For example, none have yet demonstrated fully volumetric millimeter-scale imaging depths, despite such depths being theoretically within the scattering limit of most biological tissues. Unfortunately, robustly characterizing these inverse-scattering methods is significantly challenging, largely because accounting for multiple scattering prevents an explicit analytical relationship between the measured scattering data and the sample’s 3D RI. As a result, these methods are often formulated as computational inverse problems, solved via non-convex optimization. In such cases, 3D RI is often reconstructed by iteratively adjusting RI values of each voxel within the sample volume, with the goal of minimizing the difference between scattering measurements and predictions made by the scattering model. Importantly, there is no mathematical guarantee that this strategy yields a converged solution that is physiologically true or even realistic. Indeed, it is well-known in other fields that the converged solution to a non-convex inverse problem is highly sensitive to factors such as the quality and quantity of input data, the choice of optimization algorithm, the conditions under which convergence is performed, the incorporation of appropriate computational priors, or even the complexity of the true solution itself^60-63^. To the best of our knowledge, this issue has not yet been explored in the context of inverse-scattering in the optical imaging realm.

In this work, we investigate how various computational factors influence the converged 3D RI reconstruction using the multi-slice beam propagation (MSBP) inverse-scattering method. MSBP is a relatively simple yet surprisingly effective inverse-scattering method^36,49,53,64-68^. However, as we will show in this manuscript, MSBP suffers from local minima similar to most other nonconvex inverse-problems. Specifically, we will show that the quality of MSBP-based 3D RI reconstruction is critically, and sometimes counter-intuitively, affected by the seemingly arbitrary choice of whether to use a complex-valued field versus real-valued amplitude cost function, computational zero-padding, Rytov-based initial guess, or optical defocusing. Additionally, we will outline the methodology that we empirically found to be most effective for MSBP to robustly reconstruct high-quality RI distributions across diverse biological samples. All code and data used to generate these results are open source so that others can replicate our work.

Our hope is to stimulate follow-up works that explore best practices with other more sophisticated inverse-scattering models, which may exhibit different convergence characteristics. Ultimately, our long-term goal is to advance the field of inverse scattering to enable computational scatter-correction and deep-tissue image reconstruction of thick multicellular samples, overcoming the current limitations of conventional techniques for large-scale, volumetric biological imaging.

## Results

### Basic principles of computational optics

We begin by highlighting three computational optics techniques that form the foundation of our approach to robustly conduct inverse scattering in various multiple-scattering samples.

#### Computational propagation of fields

Propagation of an optical field through homogenous media can be efficiently performed computationally using principles from Fourier optics, which treat the field as a superposition of plane waves with different spatial frequencies^69^. The process begins by taking a two-dimensional spatial Fourier transform of the complex-valued field distribution at an initial plane, which decomposes it into its component plane waves (i.e., angular spectrum). Each plane wave component corresponds to a specific spatial frequency and propagation direction.

To propagate the field over a distance Δ*z*, each plane wave is multiplied by a complex phase factor known as the transfer function of free space, which accounts for the phase shift accumulated by that corresponding wave during propagation. This transfer function depends on the spatial frequencies and the wavelength, typically expressed as,

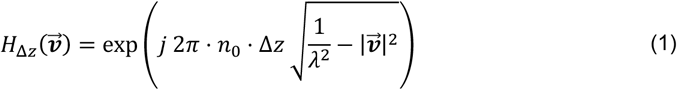

where 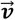 is the 2D transverse (*v*_*x*_, *v*_*y*_) spatial-frequency of the component plane wave, *λ* is the wavelength, and *n*_0_ is the RI of the homogenous media that the wave is propagating through. After applying this phase factor to each component plane wave, an inverse Fourier transform is performed to reconstruct the field after being propagated by distance Δ*z*.

### Inverse scattering based on weak-scattering approximation and angular scattering field measurements

For weakly scattering samples, the Fourier diffraction theorem provides a crucial relationship between the measured scattered fields and the sample’s 3D RI distribution in spatial-frequency space (k-space)^70,71^. Specifically, under plane wave illumination from a given direction, the measured scattered field maps to a spherical shell in the sample’s k-space, with the shell’s position determined by the incident and scattered wave vectors. Thus, each illumination angle maps sample-specific information to unique spherical shells in k-space.

By systematically varying the illumination angle and recording the resulting fields, different regions of k-space are sampled. As more illumination angles are used, these spherical shells accumulate, gradually filling k-space and enabling reconstruction of the full 3D RI distribution via inverse Fourier transform. This multi-angle acquisition strategy is essential for accurate, high-resolution 3D RI imaging, as it allows the *volumetric* reconstruction of the object’s spatial frequency distribution from a series of 2D field measurements

### Inverse scattering with multi-slice beam-propagation

At the core of our investigation is the multi-slice beam propagation (MSBP) method, which has recently shown strong potential for reconstructing 3D RI distributions in multiple-scattering regimes. MSBP models light–sample interactions by decomposing the 3D volume into a series of infinitesimally thin 2D layers, each acting as a phase-modulating mask. The sample’s bulk scattering characteristics are modeled by accumulating diffraction effects as light propagates layer-by-layer through the sample.

This light propagation process through the sample consists of two main steps: 1) wave-propagation through homogenous medium between adjacent 2D layers; and 2) phase-modulation by the layer’s complex transmittance function. These steps are mathematically expressed as shown below:

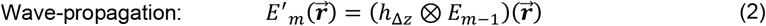

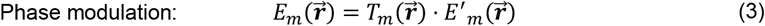

Here, 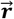 is the 2D traverse (*x, y*) spatial-coordinates variable. *h*_Δ*z*_ denotes the optical propagation kernel by the distance Δ*z* separating the layers and is the inverse Fourier transform of Eq. (1). The operator ⊗ represents mathematical convolution. *E*_*m*_ and 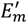 denote the *outgoing* and *incoming* 2D electric fields at the *m*^th^ layer of the 3D sample, respectively. This *m*^th^ sample layer is modeled as a phase-mask with the 2D complex transmittance function 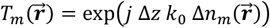, where *k*_0_ = 2*πn*_0_/*λ* denotes wavenumber, and 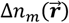 denotes the RI difference between the *m*^th^ layer of the sample and the surrounding media with homogenous RI, i.e., 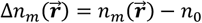.

Eq. (2) describes that the field *incoming* to the sample’s *m*^th^ layer is simply the *outgoing* field from the previous (*m* − 1)^th^ layer, propagated by the distance Δ*z* between layers. Eq. (3) computes the *outgoing* field from the *m* ^th^ layer by simply multiplying the incoming field with the layer’s transmittance function. Thus, by repeatedly implementing Eqs. (2) and (3), an illumination wave incident to the sample can be fully propagated through all sample layers before being measured. This constitutes the MSBP forward model, which takes as input a 3D RI distribution as well as an incident wave, and computes the scattered field after the incident wave propagates all the way through. This sequence defines the MSBP *forward model*, which predicts the scattering measurement from a given 3D RI distribution and incident illumination. The goal of inverse-scattering is to “invert” this forward model and reconstruct the sample’s RI from collected measurements. This is often implemented as a least squares problem that iteratively searches for a solution to the sample’s 3D RI distribution that minimizes the difference between experimental measurements and model predictions:

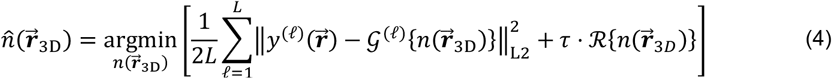

Here,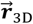 denotes the 3D spatial-coordinate (*x, y, m*), where *m* indexes the axial layers of the sample, as illustrated in **Figure 1**, i.e., 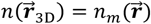 for *m* = 1,2, … *M* for a *M*-layer sample. The MSBP forward model is represented by the operator *𝒢*, which encapsulates the repeated application of Eqs.(2) and (3), based on some estimate of 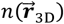. The raw scattering measurement is denoted by 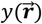, with the parameter *ℓ* indexing the measurement (described below). Lastly, ‖… ‖_L2_ denotes the L2-norm and *ℛ*{.} represents a regularization operator to stabilize convergence. While various regularization strategies are available, in this work we use total-variation. The parameter *τ* controls the strength of regularization and is empirically tuned. We solve the minimization-based inverse problem in Eq. (4) using gradient descent, which also requires an empirically selected step size. To identify an effective combination of step size and regularization strength, we performed a series of MSBP-based inverse-scattering RI reconstructions on calibration microspheres with known refractive indices. We selected the parameter pair that yielded quantitatively accurate RI reconstructions for these reference spheres (see Supplementary Information).

**Figure 1.**
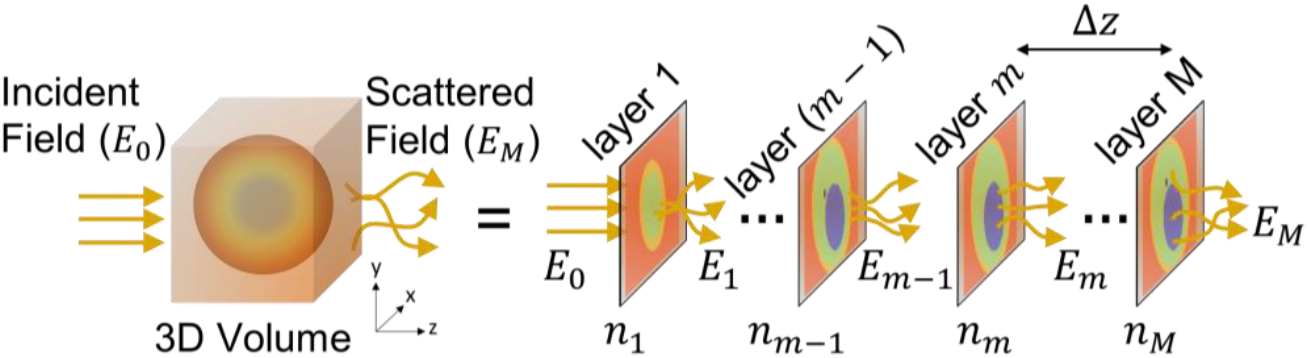
Schematic of MSBP forward model. MSBP forward model treats 3D object as multiple thin 2D transmission layers and scattered field is modeled via sequential layer-to-layer electric-field propagation through the sample.

Importantly, reconstructing the sample’s 3D RI distribution is the primary objective of inverse scattering. *If successfully reconstructed, this RI distribution can be directly interpreted as a volumetric image of the sample that provides label-free and quantitative contrast of its endogenous morphology*.

### Strategies for MSBP-based inverse-scattering

Due to the inherent complexity of light scattering, solving for 3D RI from 2D measurements using Eq. (4) is an ill-posed problem. To improve its conditioning and enable more reliable reconstruction, it is essential to introduce diversity in the measured data. Below, we detail our data acquisition and pre-processing strategies designed to enhance measurement diversity, along with empirical observations into which approaches most effectively stabilized convergence.

#### 1. Field versus amplitude-only measurements

The scattering measurement 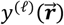 can be acquired either as complex-valued field or real-valued amplitude (i.e., square-root of intensity) measurements. The forward model must be adapted accordingly. Let 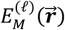 denote the field exiting the final (*M*-th) layer of an *M*-layer sample. In the case when directly using complex-valued field measurements (neglecting defocusing and resolution filtering) in the Eq. (4) inverse solver, the forward model corresponds directly to this field: 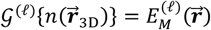. In contrast, when using amplitude-only measurements in Eq. (4), the forward model returns the *magnitude* of the field, i.e., 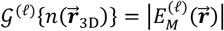.

#### 2. Measurements with angled illumination and sample defocus

Diversity in the scattering measurements can be achieved by illuminating the sample at different angles and/or axially defocusing the sample. In the case of angular plane-wave illumination, the field *before* the sample’s first layer can be expressed as 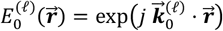, where 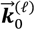 denotes the lateral wavevector of the plane-wave corresponding to the *ℓ*-th illumination angle. The forward model naturally accounts for this by using Eqs. (2) and (3) to propagate 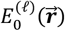 through the sample. To model sample defocus, an additional propagation step (i.e., Eq. (1)) can be applied to the field exiting the sample, effectively shifting the sample to the desired imaging plane.

#### 3. Initialization with weak-scattering approximation

Reconstructing the sample’s RI from Eq. (4) involves iteratively updating voxel-wise RI values within the sample’s volume until convergence. This process requires an initial RI estimate. Due to the nonconvex nature of the scattering problem, the final solution may depend on this initialization. Prior works often initialize MSBP with a uniform RI equal to the background medium. Here, we compare that approach to one using a weak-scattering estimate derived via the Fourier diffraction theorem, followed by MSBP-based refinement. Our strategy for weak-scattering initialization employs the Rytov approximation.

To acquire diverse scattering measurements, we use an interferometric system with angle-scanning capabilities that allow precise tilting of the illumination onto the sample (see **Figure *2***). The scattered light is interfered with an off-axis reference beam, and standard digital filtering isolates the complex-valued scattered field (amplitude and phase) in *k*-space^72^. This process is repeated with the sample translated out of view in order to collect background field measurements. Sample field measurements can then be normalized by complex-dividing with the background fields. Amplitude-only “measurements” can be synthetically generated by simply taking the amplitude component of the complex-valued field measurement.

**Figure 2.**
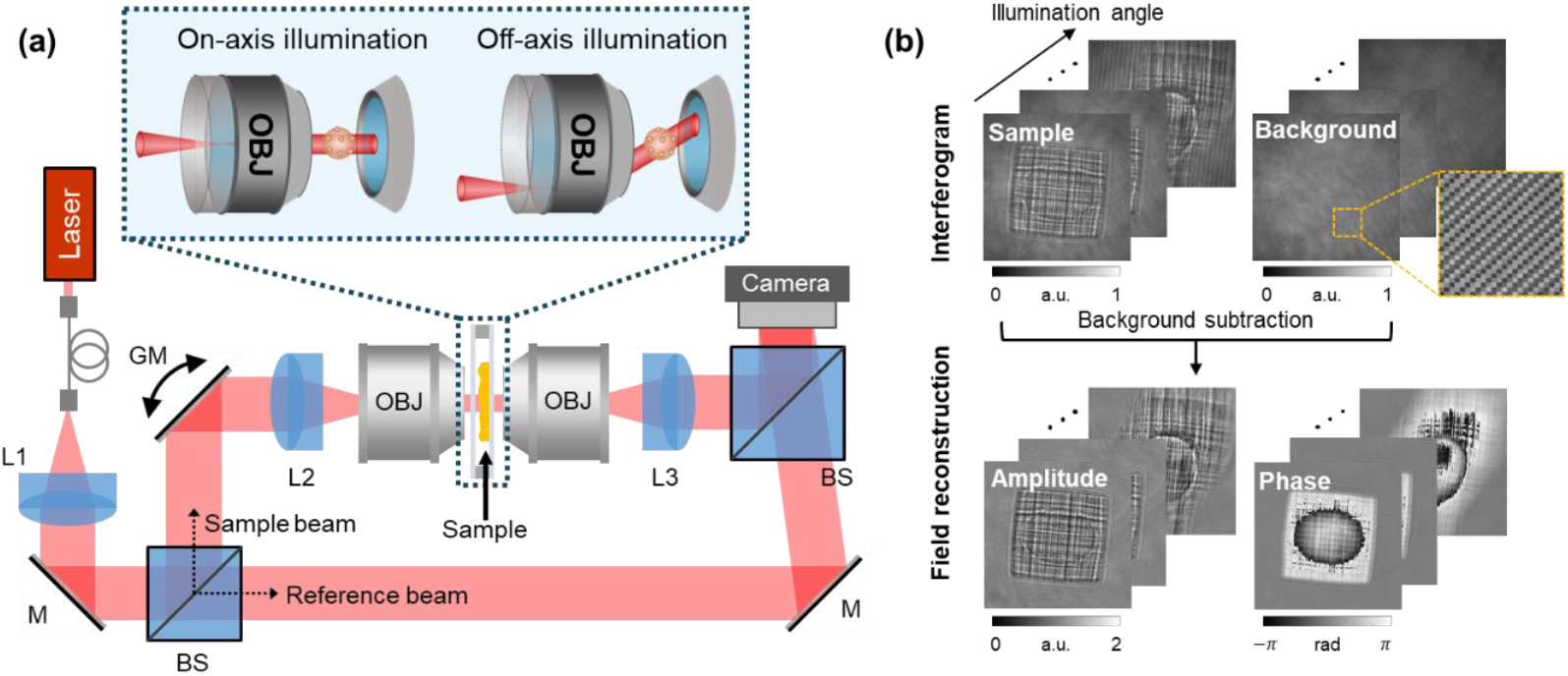
Schematic of the experimental setup and data acquisition framework. **(a)** Mach–Zehnder off-axis interferometer system with angularly scanned illumination. (L: lens, M: mirror, BS: beam splitter, GM: galvo mirror, OBJ: objective lens). **(b)** Example of data acquisition using a microphantom. Interferograms of the sample and background are sequentially captured at multiple illumination angles, followed by reconstruction of 2D amplitude and phase maps of the sample for each angle.

The normalized angular scattering field measurements can be used with the Fourier diffraction theorem to estimate the sample’s 3D RI under the weak-scattering approximation. To bypass this approximation, MSBP can be employed to reconstruct RI using either complex-valued field data or amplitude-only measurements. Additionally, digital defocus can be introduced by computationally propagating the measured field, effectively allowing for RI reconstruction at various focal planes. *As a result, field-based, amplitude-only, and defocus-varied reconstructions can all be performed from a single measurement dataset without having to physically re-acquire images*. This provides a fair and consistent basis for comparing the different reconstruction strategies that we explore below.

### Inverse-scattering with complex-valued field versus amplitude-only measurements

We begin by comparing the reconstruction performance of MSBP-based inverse scattering using either complex-valued field data or amplitude-only measurements. Note that this choice results in slight modifications to the cost function as described previously, thereby altering the inverse problem in Eq. (4). We will show that this slight modification yields dramatic changes in 3D RI reconstruction.

We note that previous studies have performed MSBP-based 3D RI reconstruction using either complex-valued field data^64,73^ or amplitude-only measurements^49,50^. However, these two approaches have not been directly compared on a multiple-scattering sample under identical experimental conditions. As a result, it has remained unclear under which circumstances one strategy may outperform the other. In this section, we present the first side-by-side comparison of both methods across real-world samples with varying scattering properties. This analysis allows us to establish a practical framework for achieving robust RI reconstruction across a range of scattering samples.

#### Comparison of MSBP-based inverse-scattering strategies with 3D scattering microphantoms

##### Phantom

We first conduct these comparison experiments using 3D scattering phantoms consisting of a 3D cell imaging target embedded into scattering cubes of increasing size. Phantoms were 3D-printed (further described in Methods) and thus internal microstructures have known geometry and 3D RI distribution^74^. **Figure 3** shows the cell imaging target, which contains internal test structures of known RI distributed in 3D, with the smallest feature size measuring 300 nm. The surrounding scattering cubes are composed of pseudo-randomly distributed rods designed to induce multiple scattering. By varying the cube size (40, 60, 80, and 100 µm), the scattering strength of the phantoms can be systematically controlled; i.e., the larger the cube, the greater the degree of multiple-scattering. This makes the phantoms a robust platform for characterizing and identifying artifacts in 3D RI reconstructions under varying levels of multiple scattering. As described previously, complex-valued field measurements of the phantoms were acquired interferometrically under varying illumination angles. Corresponding amplitude-only measurements were synthetically derived by extracting the magnitude component from the complex field data.

**Figure 3.**
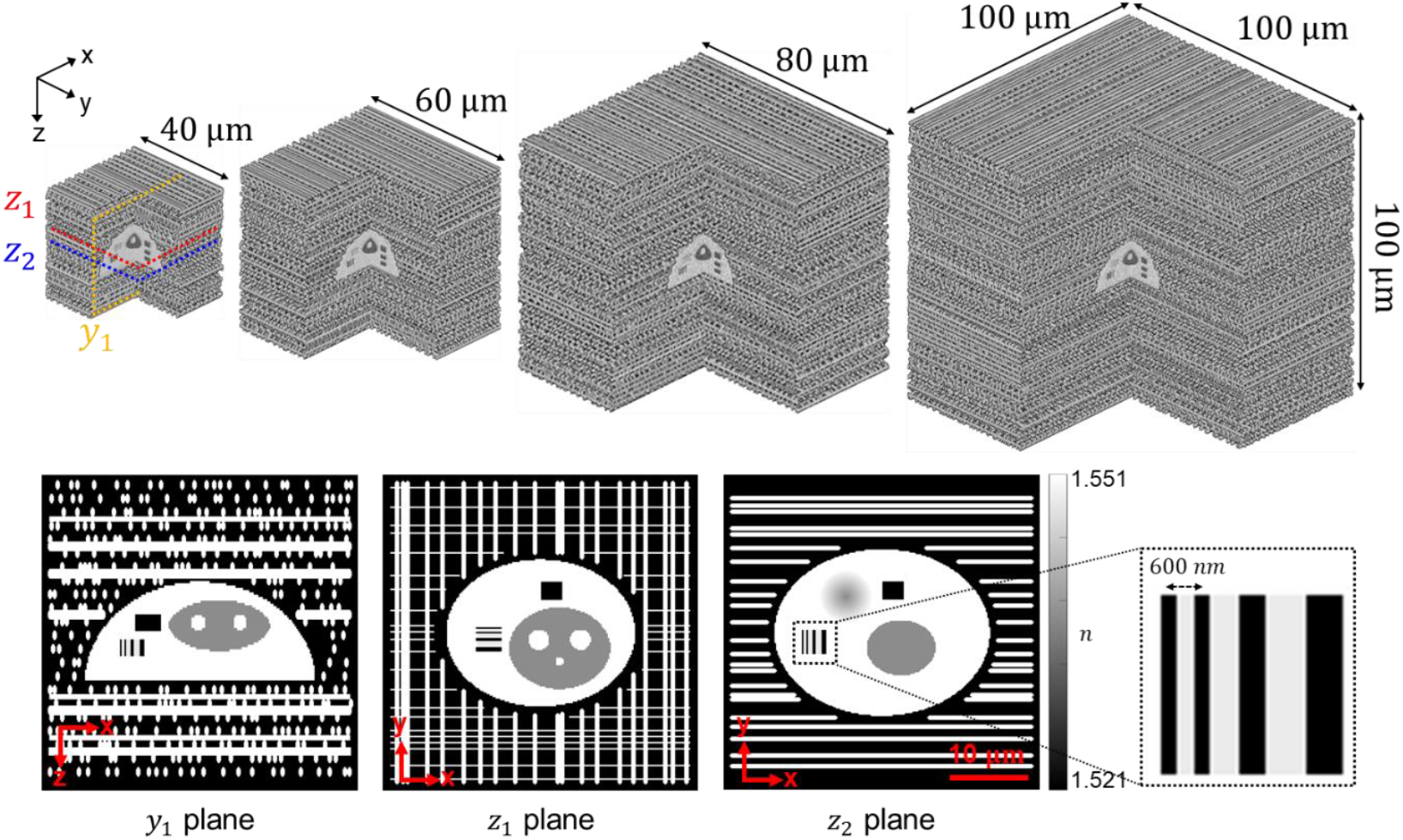
Design of a 3D scattering phantom for controlled multiple scattering. The phantom features an internal test structure embedded within a pseudo-random array of rods engineered to scatter light and obfuscate the imaging target. The extent of multiple scattering is tunable by varying the dimensions of the surrounding scattering volume.

##### Imaging results

**Figure 4(a1–a4)** present the 3D RI reconstruction results for the 40, 60, 80, and 100 µm scattering phantoms using MSBP-based inverse scattering with amplitude-only data. To output these reconstructions, the inverse-solver was initiated with a constant estimate of the sample’s 3D RI equal to that of the background medium (i.e., Δ*n* =0). As the phantom size increased, resolving fine features became more difficult; however, the overall structures were recovered without prominent artifacts.

**Figure 4.**
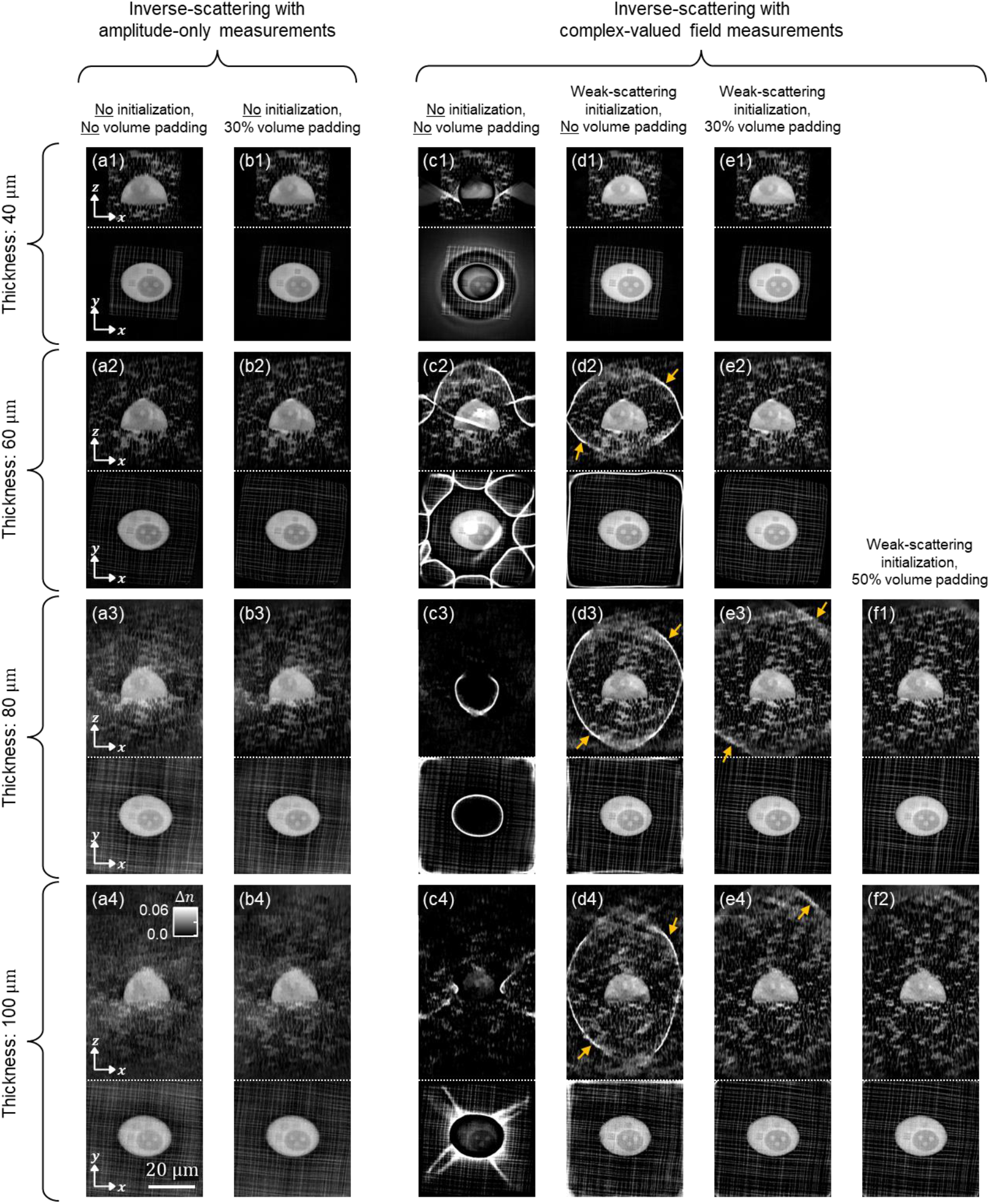
Comparison of 3D RI reconstruction results for 40, 60, 80, and 100 µm scattering cube phantoms, using amplitude-only data versus complex-valued field measurements within the MSBP inverse solver framework. For amplitude-only data, the solver was initialized with a uniform RI distribution (*Δn* = 0). 3D RI reconstruction results are shown **(a1–a4)** with and **(b1–b4)** without 30% volume padding. With complex-valued field measurements, 3D reconstruction results are shown with **(c1-c4)** *Δn* = 0 RI initialization without padding, **(d1-d4)** RI initialization with weak-scattering approximation without padding, **(e1-e4)** weak-scattering initialization + 30% volume padding, and **(f1, f2)** weak-scattering initialization + 50% volume padding. Yellow arrows in (d2-d4) and (e2, e3) correspond to reconstruction artifacts from limited FOV.

In contrast, **Figure 4(c1–c4)** show the corresponding 3D RI reconstructions using complex-valued field measurements, where the inverse-solver was also initialized with a Δ*n* =0 RI distribution. In this case, reconstruction performance deteriorates significantly, even for the smallest phantom. For example, ringing artifacts surrounding the imaging target appear in the 40, 80, and 100 µm cases, and additional spurious artifacts are observed spread across the reconstruction volume, particularly for the 60 and 100 µm phantoms. Visual inspection of the phase maps in the measurements (see **Figure 2(b)** and Supplementary Information) reveals a strong spatial correlation between phase-wrapping artifacts and the ringing artifacts observed in the RI reconstructions. This relationship implies that phase wrapping introduces ambiguities in the measured phase, which can mislead the inverse solver and drive the reconstruction toward incorrect solutions.

To guide the reconstruction toward a more accurate solution, we next initialized the inverse solver with an estimate of the sample derived from the weak-scattering approximation. Specifically, we used the field measurements to compute a 3D RI estimate via the Fourier diffraction theorem, which then served as the initial guess for the iteration-based MSBP reconstruction. The resulting RI reconstructions are shown in **Figure 4(d1–d4)**. As expected, this warm initialization led to substantial improvements. In the 40 µm scattering phantom, the reconstruction is virtually artifact-free. While artifacts persist in the larger phantoms, they are markedly reduced compared to those in **Figure 4(c1– c4)** and are no longer confined to regions near the imaging target, suggesting that a well-chosen initialization effectively mitigates ambiguities introduced by phase wrapping.

Notably, the remaining artifacts in **Figure 4(d1–d4)**, as indicated by yellow arrows, appear predominantly at the edges of the imaging field-of-view (FOV). This suggests that they are not intrinsic to the sample itself but rather a consequence of FOV limitations. This is consistent with the behavior of high-angle illumination, which can shift spatial information from sufficiently large samples to be partially outside the detectable FOV, thus resulting in missing information. This likely explains why the 40 µm phantom in **Figure 4(d1)** shows no edge artifacts, while the larger phantoms do. To test this hypothesis, we zero-padded the reconstruction volume to mitigate edge-related artifacts, while still keeping the weak-scattering initialization. **Figure 4(e1–e4)** show the results with 30% additional padding – the 40 and 60 µm phantoms are virtually artifact-free, while the edge artifacts in the 80 and 100 µm reconstructions (indicated by yellow arrows) show substantial improvement. Increasing the padding to 50% (**Figure 4(f1, f2)**) effectively eliminates the remaining edge artifacts, yielding visually artifact-free images for all phantom samples. Interestingly, as compared between **Figure 4(a1–a4)** and **Figure 4(b1–b4)**, MSBP-based inverse scattering using amplitude-only measurements did not exhibit any noticeable edge artifacts and was not significantly affected by padding.

In summary, we empirically found that image clarity was slightly higher when solving the MSBP inverse problem using complex-valued field data compared to amplitude-only measurements. However, this improvement was only realized when the field data was combined with sufficient volume padding and a warm initial guess. Without these preprocessing steps, the reconstructions were degraded by severe artifacts. For our phantom sample, a weak-scattering approximation provided a sufficiently good initial guess. However, biological specimens may exhibit more complex scattering properties, making the weak-scattering approximation inadequate. Additionally, larger samples may require extensive padding, which can significantly increase computational cost. In contrast, reconstructions using amplitude-only measurements required neither an initial guess nor volume padding to produce comparable image quality, thus offering a more computationally efficient approach less dependent on prior-knowledge. We next investigate whether these trends persist in real-world biological specimens.

##### Comparison of MSBP-based inverse-scattering strategies with 3D scattering biological sample

We next examine whether the MSBP-based inverse-scattering reconstruction strategies demonstrated in phantoms translate effectively to a real-world biological sample. Here, we use an intestinal organoid as an example of a complex 3D scattering specimen. Guided by insights from our phantom experiments, we first implemented the MSBP inverse-scattering solver using complex-valued field measurements acquired at multiple illumination angles. To avoid edge-related artifacts, we implemented sufficient volume padding equal to 100% of the imaging field-of-view.

As a baseline, the inverse solver was first initialized with a uniform RI distribution equal to the background medium (Δ*n* =0). As shown in **Figure 5(a1–a4)**, this approach resulted in pronounced artifacts distributed throughout the organoid’s 3D reconstruction. This outcome mirrors the artifacts observed previously in **Figure 4(c1–c4)**, where the phantom data underwent the same reconstruction conditions. This suggests that similar challenges arise in both synthetic and biological scattering scenarios when the reconstruction is poorly initialized and complex-valued field measurements are used in the inverse-scattering solver.

**Figure 5.**
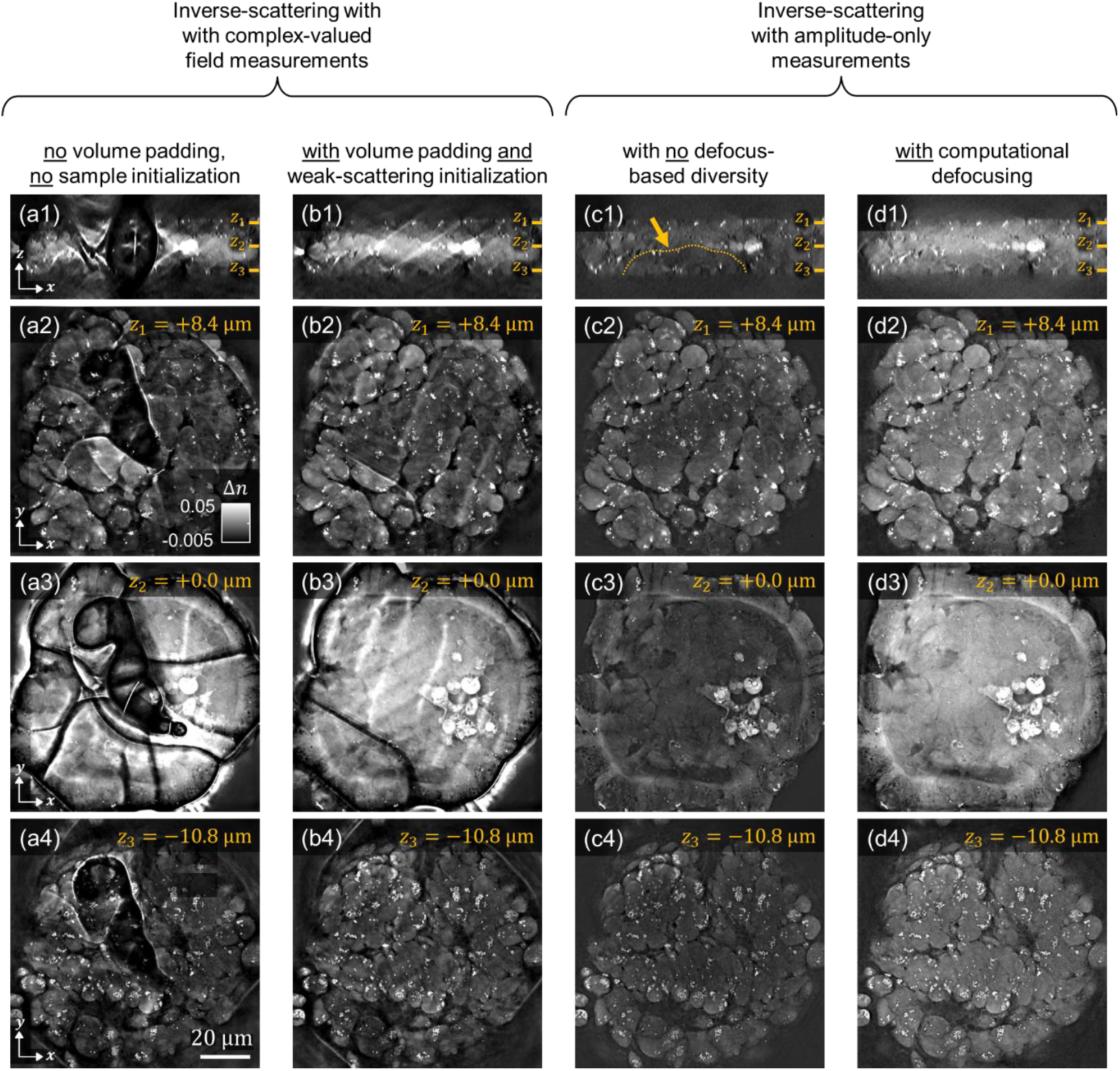
Comparison of MSBP-based inverse-scattering strategies for 3D RI reconstruction of an intestinal organoid sample. Axial cross-sections are shown from the 3D RI reconstructions obtained using different inverse-solving strategies: **(a1)** reconstruction using complex-valued field measurements with a uniform initial guess (*Δn* =0); **(b1)** reconstruction using complex-valued field measurements with an initial guess derived from the weak-scattering approximation; **(c1)** reconstruction using amplitude-only measurements without any initialization, and **(d1)** reconstruction using amplitude-only measurements with additional defocused measurements to introduce depth diversity. For each axial cross-section, corresponding RI reconstructions at two lateral planes are also shown. Specifically, **(a2, a3), (b2, b3), (c2, c3)**, and **(d2, d3)** display lateral cross-sections located at z = –4.2 µm and +6.6 µm relative to the axial planes shown in **(a1), (b1), (c1)**, and **(d1)**, respectively.

We then initialized the inverse solver with a 3D RI estimate derived from the weak-scattering approximation. In our earlier phantom experiments, this strategy of using complex-valued field measurements combined with a weak-scattering-based initial guess proved effective in driving convergence to high-quality RI reconstructions. Interestingly, we observe a different outcome here. As shown in **Figure 5(b1–b4)**, while the weak-scattering initialization does lead to improved results compared to the constant Δ*n* =0 initial guess, the final 3D reconstruction still contains pronounced artifacts throughout the volume (indicated with yellow arrows). This suggests fundamental differences in the scattering behavior of the organoid versus the phantom. Specifically, the organoid appears to exhibit significantly stronger multiple scattering, rendering the weak-scattering approximation insufficient as a warm start for the inverse solver. This interpretation is supported by a comparison of raw phase measurements (see Supplementary Information, **Figure S2**), where the organoid displays substantially more chaotic phase-wrapping than the phantom, indicating greater optical heterogeneity and a higher degree of multiple scattering.

We next implemented the inverse-scattering solver using amplitude-only measurements, *without* employing an initial guess or volume padding. As shown in our earlier phantom experiments, this strategy demonstrated a surprising robustness to reconstruction artifacts across a range of scattering strengths, while still achieving reconstruction quality comparable to that of field-based measurements when combined with warm initialization and volume padding. Consistent with those findings, the results shown in **Figure 5(c1–c4)** exhibit similar behavior here. Unlike the field-based reconstructions in **Figure 5(a1–a4)** and **Figure 5(b1–b4)**, no sharp spatial discontinuities or artifacts are observed. Structural features such as lipid droplets, apoptotic cells, and cellular boundaries can be clearly resolved.

However, one notable artifact is the presence of a low-contrast cavity region within the 3D organoid, which does not align with the expected physiological morphology. This region appears axially localized and lies beneath highly scattering structures such as apoptotic cells and lipid aggregates. We speculate that these features may scatter light away from the underlying cavity, misleading the inverse solver to converge to a solution containing a “shadow” beneath these high-RI structures (see dashed yellow line in **Figure 5(c1)**). Similar shadowing artifacts are well-documented in other scattering-based imaging modalities, such as optical coherence tomography (OCT)^75^.

To mitigate this effect, we introduced defocus diversity into the reconstruction process to increase axial sensitivity and stabilize reconstruction of features that may otherwise be occluded by high-RI structures. We do this by synthetically generating amplitude-only measurements from multiple defocused planes. This was achieved by computationally propagating the raw field measurements and extracting their amplitude components. As shown in **Figure 5(d1–d3)**, incorporating these defocused measurements significantly improves reconstruction quality by eliminating the shadow artifact and restoring contrast in the cavity region, thereby enabling clear visualization of the full organoid volume. This approach thus overcomes the major limitation of amplitude-only reconstruction without defocus diversity, which is that the sample’s phase information is disregarded. By leveraging the phase information indirectly encoded within defocused measurements, but not directly using the phase within the inverse-solver to avoid phase-wrap artifacts, the inverse-scattering reconstruction process becomes more robust and yields more accurate representations of internal structures.

By combining insights from both the phantom and organoid experiments, we observe a consistent trend: inputting complex-valued field data from multiple illumination angles into the inverse solver without warm initialization results in 3D reconstructions corrupted by sharp, discontinuous artifacts when imaging multiple-scattering samples. Initializing the solver with a 3D RI estimate from the sample’s weak-scattering approximation while still using complex-field data in the solver helps stabilize the reconstruction and may be sufficient when the sample exhibits only mild multiple scattering, as seen with the phantom. However, for more strongly scattering samples like the organoid, this weak-scattering initialization is inadequate, and significant artifacts persist. In contrast, amplitude-only measurements across multiple illumination angles proved to be a more robust strategy, producing artifact-free reconstructions in both phantoms and organoids *without* requiring any prior initialization. Nonetheless, even this approach may exhibit shadowing artifacts beneath highly scattering structures, reducing image clarity in deeper regions. To address this, we introduced defocus diversity by incorporating synthetically defocused amplitude-only measurements. This strategy of combining angular and defocus diversity significantly improved reconstruction quality and eliminated axial shadowing, enabling more uniform and high-clarity visualization of biological morphology throughout the organoid, as well as other biologically scattering samples (see Supplementary Information). With this methodology established, we now apply it to additional biologically relevant scattering samples to evaluate its broader utility.

### Biological results

#### Mutant C. elegans strain with increased lipid synthesis

We first demonstrate the capability of this inverse-scattering methodology by reconstructing the 3D RI of a *daf-2(e1370)* mutant *C. elegans* worm. The *daf-2(e1370)* mutant of *C. elegans* is a temperature sensitive partial loss of function mutant with elevated lipid synthesis and storage compared to the wild type due to decreased insulin/IGF-1 signaling^76,77^. The abundant internal lipid content in *daf-2(e1370)* mutants lead to increased optical scattering due to their different RI compared to the surrounding tissues^78^. **Figure 6(a1)** presents a lateral cross-sectional view of the central XY plane from the 3D image reconstructed via the methodology described previously. Despite the strong scattering caused by the high lipid concentration, the 3D structural features of the *C. elegans* were successfully recovered with remarkably high contrast. A magnified region is also shown, highlighting an area containing *C. elegans* embryos. **Figure 6(a2)** presents a zoomed-in view of the worm’s embryo region, clearly revealing anatomical structures such as the intestinal lumen and vulva. Further magnified views in **Figure 6(a3, a4)** show individual embryos at subcellular resolution, with distinct cell boundaries and nuclei visible. Notably, the embryos display different cell counts that mark their respective developmental stages. This level of detail can be particularly valuable when studying embryonic development, especially when individual embryonic cells and nuclei can be volumetrically visualized *within the intact worm*. Combined with its capabilities for label-free morphological contrast, this imaging method thus provides a powerful tool for potentially identifying and analyzing different stages of cell development *in-vivo*.

**Figure 6.**
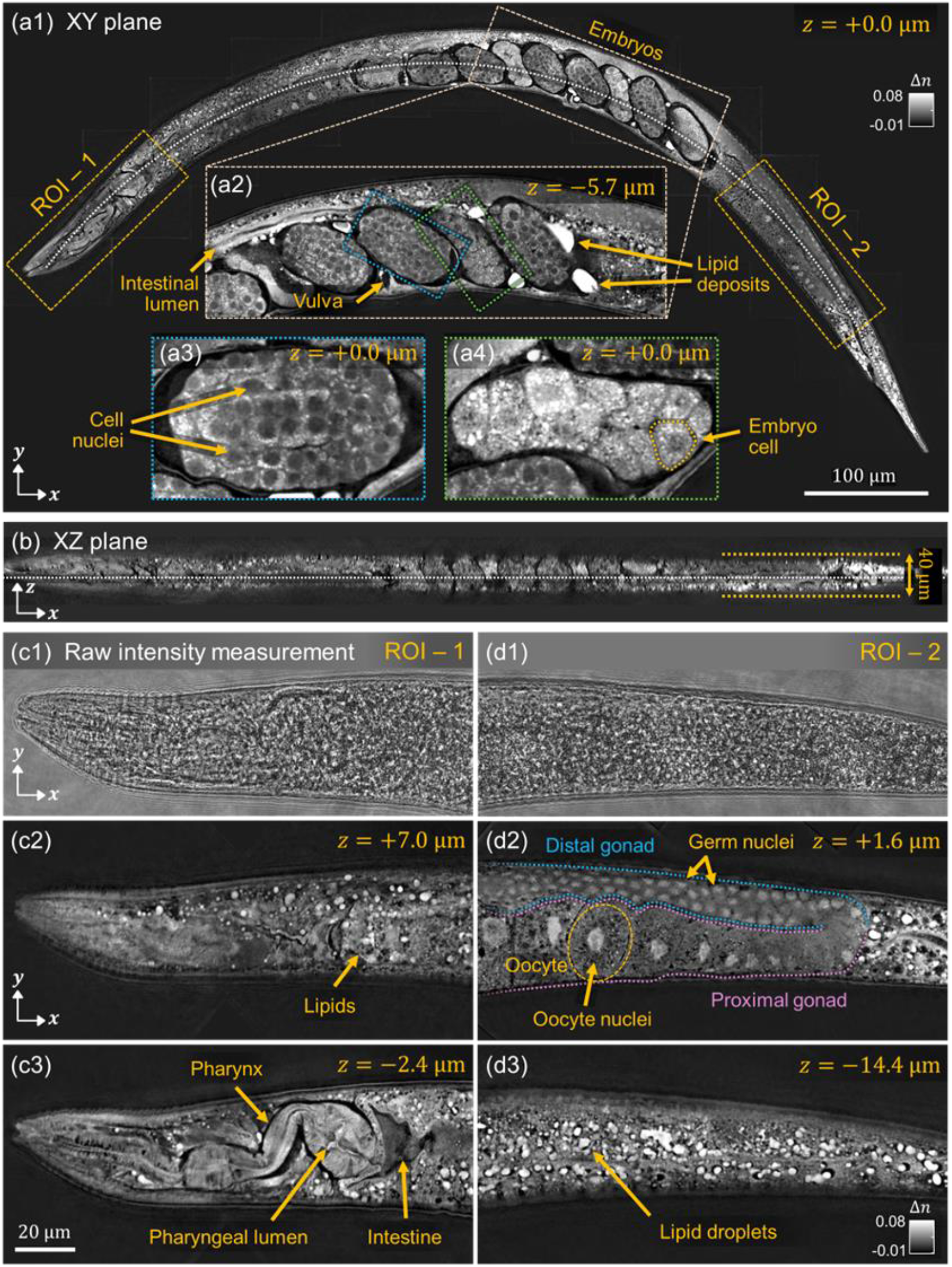
RI reconstruction results obtained in *daf-2(e1370)* mutant *C. elegans* samples, which is characterized by increased lipid accumulation. **(a1)** Cross-sectional view is shown of the central XY plane, with various structures such as embryos, intestinal lumen and vulva. (**a2**) A zoomed-in image is shown of the region with embryos at different development stages. **(a3, a4)** Further magnified views of individual embryos show subcellular resolution, with embryonic cell boundaries and nuclei clearly visible. **(b)** Volumetric imaging capability is demonstrated by showing a curvature-corrected XZ plane along the dashed white line in (a). **(c1-c3, d1-c3)** Raw scattering measurements (1) and corresponding 2D cross-sectional views are shown for two regions of interest (ROI) (2,3). ROI-1 includes the head and pharynx region, and ROI-2 covers the tail region; both marked by boxes in the cross-sectional view in (a). The intestine, lipids, pharyngeal lumen and pharynx in ROI-1, as well as the proximal and distal gonads, germ nuclei, oocytes, oocyte nuclei, distal gonad and lipid droplets in ROI-2, are indicated by arrows.

To highlight 3D structure, **Figure 6(b)** shows a curvature-corrected axial cross-sectional XZ plane along the dashed white line in **Figure 6(a1). Figure 6(c1-c3) and 6(d1-d3)** shows representative raw measurements and the corresponding reconstruction results from two different regions-of-interest (ROI) and imaging depths in the mutant *C. elegans*. ROI-1 corresponds to the anterior head region, while ROI-2 includes the tail region. In **Figure 6(c1, d1)**, the raw intensity images captured under standard illumination show strongly scattered light, which obscures internal features due to the complex scattering within the sample. In contrast, the rich RI-based label-free contrast from the inverse scattering reconstructions in **Figure 6(c2, c3)** and **Figure 6(d2, d3)** reveal detailed internal structures across different depths, including the pharynx, pharyngeal lumen, intestine, proximal gonad, germ nuclei, oocyte, oocyte nuclei, distal gonad, and lipid-rich regions. Lipid-rich regions appear as bright, spherical, vesicle-like structures due to their elevated RI and correspondingly increased optical scattering properties^78^. Their morphology is consistent with previous studies, which showed that major fat stores in *C. elegans* are contained within distinct, spherical cellular compartments^76^. Furthermore, high-quality volumetric visualization of intact cellular structures together with individual lipid droplets provides valuable insights into internal lipid organization without the need for additional labels or dyes. This capability is especially promising for studies on lipid metabolism, metabolic disorders, and the aging process^79,80^.

Notably, these imaging results correspond to the reconstruction obtained using amplitude data *with* digital refocusing. In contrast, reconstructions based on amplitude data *without* digital refocusing (i.e., raw amplitude data collected from one imaging plane) fail to resolve the internal structures (see Supplementary Information, **Figure S3**). Specifically, while the overall outline of the organism is still visible, fine internal features remain indistinct. We note that 3D reconstruction of a wild-type *C. elegans*, which does not have elevated lipid synthesis and is thus less optically scattering, does not seem to be as heavily affected by whether digital refocusing is used (see Supplementary Information, **Figure S4**). This result demonstrates that incorporating digitally refocused data is most effective for samples with complex internal scattering.

#### Intestinal organoid

Next, we performed MSBP-based inverse-scattering of an intestinal organoid larger than the one shown previously in **Figure 5**. Organoids are three-dimensional (3D) cellular structures derived from stem cells that closely mimic the architecture and function of the tissue of origin. These models have emerged as powerful tools for studying tissue development, host-microbiome interactions, and drug-induced toxicity, with more physiological relevance than traditional 2D cultures^81-86^. However, imaging organoids presents challenges due to their multicellular organization and size, which leads to substantial light scattering. Traditional imaging systems such as brightfield and epifluorescence microscopy often lack sufficient optical sectioning and are prone to high background noise, making them suboptimal for volumetrically resolving internal features in 3D organoids^87^. While confocal and light-sheet microscopies enable optical sectioning for 3D organoid imaging, they typically rely on fluorescent labeling, which is subject to photobleaching and can disrupt organoid function^88-91^.

To address these limitations, we evaluated the performance of our inverse-scattering framework for label-free 3D RI imaging of an intestinal organoid sample with a physical thickness of approximately 180 µm. In our previous experiments, we utilized a high numerical aperture (NA) 1.42 oil immersion lens to achieve high-resolution imaging. However, the short working distance of such lenses restricted the imaging depth, limiting their applicability for the thicker organoid specimens. Therefore, we adopted a 0.8 NA water-dipping objective for this study, which offered a longer working distance that could accommodate the relatively thick size of the organoids.

**Figure 7** presents the 3D RI reconstruction results of the intestinal organoid. **Figure 7(a)** shows a 3D rendered, depth-coded tomographic view, while **Figure 7(b1, b2)** displays maximum intensity projections along the *z*- and *x*-axes of the reconstructed volume, respectively. **Figure 7(c1, c2, c3)** show lateral *xy*-plane cross-sectional views at different *z* planes, corresponding to the dashed lines indicated in **Figure 7(b2)**. **Figure 7(d1, d2)** shows axial *xz*-plane cross-sectional views at different *y* positions, corresponding to the dashed lines in **Figure 7(c1)**. Together, these views volumetrically reveal important morphological features of the organoid, such as the crypts, lumen, and epithelial lining. Additionally, **Figure 7(e1, e2)** illustrate zoomed-in views that reveal densely packed cell clusters along the epithelial lining, allowing detailed subcellular visualization of cell boundaries, nuclei, lipids, and other intracellular components. Taken together, the overall morphology and the fine subcellular details revealed in the RI reconstruction offer valuable insights into the sample’s spatial organization and internal architecture.

**Figure 7.**
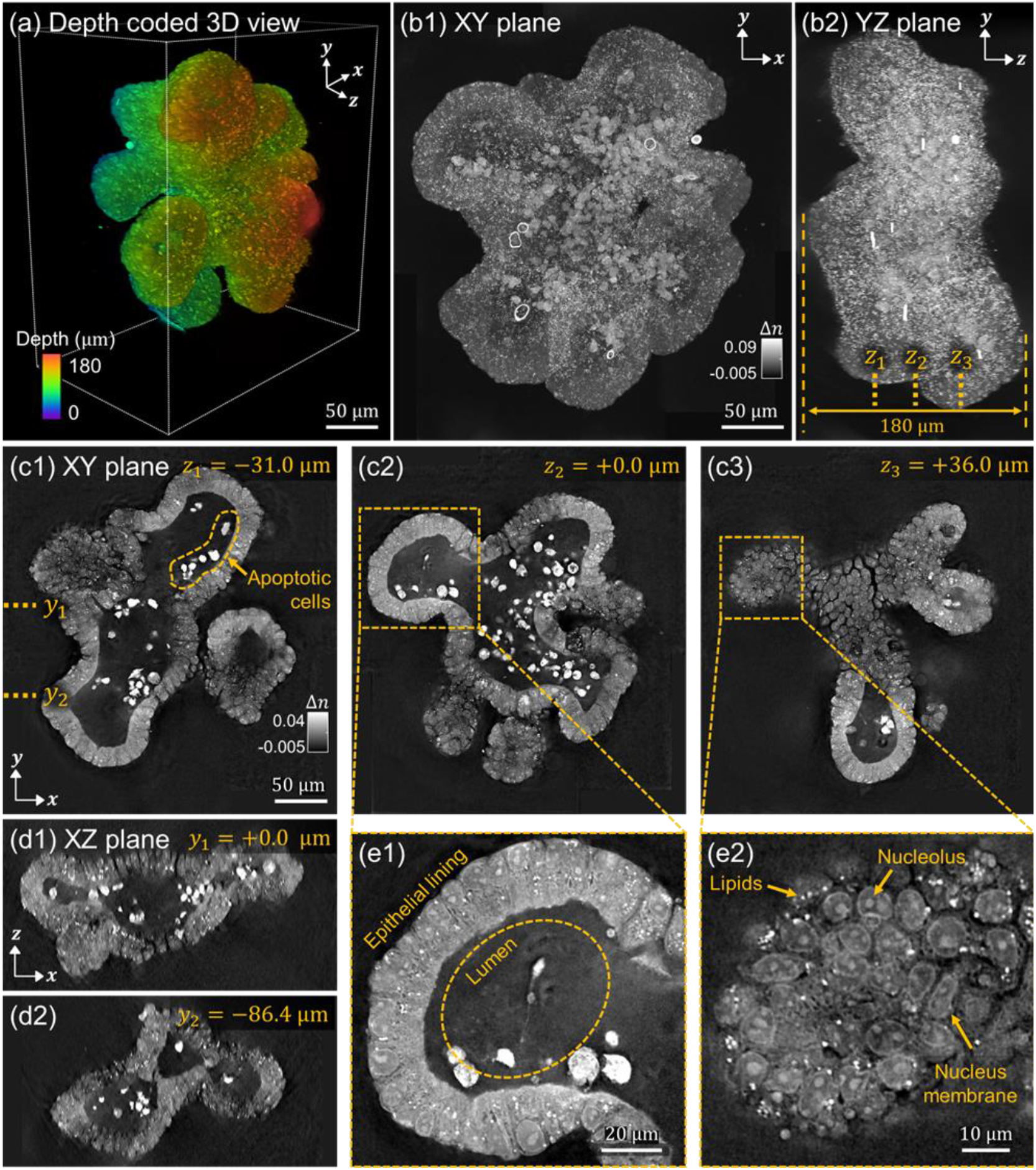
Inverse-scattering reconstruction results for intestinal organoid samples. **(a)** Depth-coded tomographic view of the reconstructed 3D RI volume. **(b1, b2)** Maximum intensity projections are shown along the *z*-axis and *x*-axis, respectively. **(c1, c2, c3)** Lateral *xy*-plane cross-sectional views are shown at different *z* planes within the reconstruction volume, corresponding to the dashed lines indicated in (b2). **(d1, d2)** Axial *xz*-plane cross-sectional views at different *y* planes of the reconstruction volume, corresponding to the dashed lines indicated in (c1). **(e1, e2)** Zoomed-in *xy*-plane lateral views of the regions outlined by dashed boxes in (c2) and (c3). Epithelial lining, lumen, lipids, nuclei, and cell boundaries are clearly visible in the images and indicated by arrows.

This imaging capability can have important applications. For example, intestinal physiology dictates a continuous cell turnover in the intestinal epithelium, with the cells at the villus tip undergoing anoikis and shedding into the lumen^92^. The presence of bright-signal (corresponding to high RI) from cell bodies within the organoid lumen is suggestive of apoptotic cells with elevated lipid content. Previous studies have consistently demonstrated that lipid accumulation occurs in epithelial cells during apoptosis^93^. This phenomenon can serve as an indication for apoptotic cell detection using magnetic resonance spectroscopy^94^. The increase in intracellular lipid content is attributed to the suppression of mitochondrial fatty acid β-oxidation, which diminishes fatty acid catabolism and enhances lipid biosynthesis during the apoptotic process^95^. Because RI-based contrast is highly sensitive to lipid content, image-reconstruction via inverse-scattering may be especially useful for the detection of apoptotic cells in certain tissues.

As before, the 3D RI reconstruction shown in **Figure *7*** was performed using amplitude-only data with computational defocusing. Without incorporating defocused measurements, sharp reconstruction artifacts appeared both around the periphery and within the organoids (see Supplementary Information, **Figure S4**). These artifacts were more severe than the slowly-varying shadow artifacts observed previously in **Figure 5** when 3D RI in a smaller intestinal organoid was reconstructed without using computational defocusing. We speculate that this is due to the specific internal structure of the intestinal organoids. Specifically, the interface between the epithelial lining and the lumen in this larger organoid constitutes a sharper RI transition that may introduce refractive effects not accurately modeled by the MSBP framework. This model mismatch may thus lead to artifacts around interfaces with abrupt RI change. By including defocused data that shifts the imaging focus away from the central lumen toward regions where the RI mismatch is less severe, these effects were observed to be significantly reduced, though not always eliminated (see Supplementary Information, **Figures S6** and **S7**).

#### Zebrafish embryo

Lastly, we show 3D RI reconstruction of a 24hpf zebrafish embryo, which is a widely used model organism in developmental biology and biomedical research. As a representative multicellular system, it provides invaluable insights into embryogenesis and organ development. However, imaging the zebrafish embryo, especially at subcellular resolution, poses significant challenges due to its complex multicellular architecture, which gives rise to substantial multiple light scattering. Accurately visualizing and quantifying its internal structures in 3D is essential for understanding developmental processes, but conventional imaging techniques often fall short due to scattering-induced degradation in resolution and contrast.

To address these limitations, we employed an MSBP-based ODT method to perform label free, high resolution 3D RI reconstruction of the zebrafish embryo. As in our imaging experiments with intestinal organoids, we used the 0.8 NA water-dipping objective lens for longer working distance. We reconstructed the sample’s 3D RI using refocused datasets, which allowed for better compensation of scattering and improved convergence during the iterative reconstruction process. This approach outperformed conventional amplitude-only MSBP-ODT reconstructions, particularly in deeper or highly scattering regions of the sample.

**Figure 8** presents detailed views from the 3D RI reconstruction, which may provide valuable insights into the spatial organization of cells and the morphogenetic processes of the embryo. For example, **Figure 8(a)** presents a central *xy*-plane slice of the reconstructed 3D RI volume, where internal anatomical features such as the spinal cord, notochord, vent, and somites are clearly visualized with subcellular resolution. The epithelial nature of the digestive track is readily evident, and a zoomed-in view clearly visualizes the vent, which is the external opening of the cloaca. **Figure 8(b)** shows a curvature-corrected XZ plane along the dashed white line in **Figure 8(a)**, clearly revealing the distribution of notochord from an axial perspective. **Figure 8(c-e)** provides magnified views of different ROIs indicated by the dashed boxes in **Figure 8(a)**. For example, **Figure 8(c1)** shows the surface of the sample, where the nuclei and cell membrane at regions of cell-cell contact between neighboring periderm cells are clearly evident. Intracellular heterogeneity, possibly due to cytoskeletal structures, are also visible. **Figure 8(c2)** shows that the nascent notochord can be visualized lower in depth, closer to the midline of the sample. Moving anteriorly along the sample, **Figure 8(d1)** shows a more developed notochord, within which prominent vacuoles are clearly distinguishable. The hypochord is visible just ventral to the notochord and the floor plate of the spinal cord is readily discernible. Imaging deeper into the sample **Figure 8(d2)**, somites are clearly visible and individual muscle fibers can be distinguished. Further anterior, **Figure 8(e1)** shows that cells within the notochord are highly vacuolated – thus, in this specific sample the maturation of the notochord can be visualized. Lower in depth, **Figure 8(e2)** shows that cells of a neuromast can be visualized situated between the muscle and surface ectoderm.

**Figure 8.**
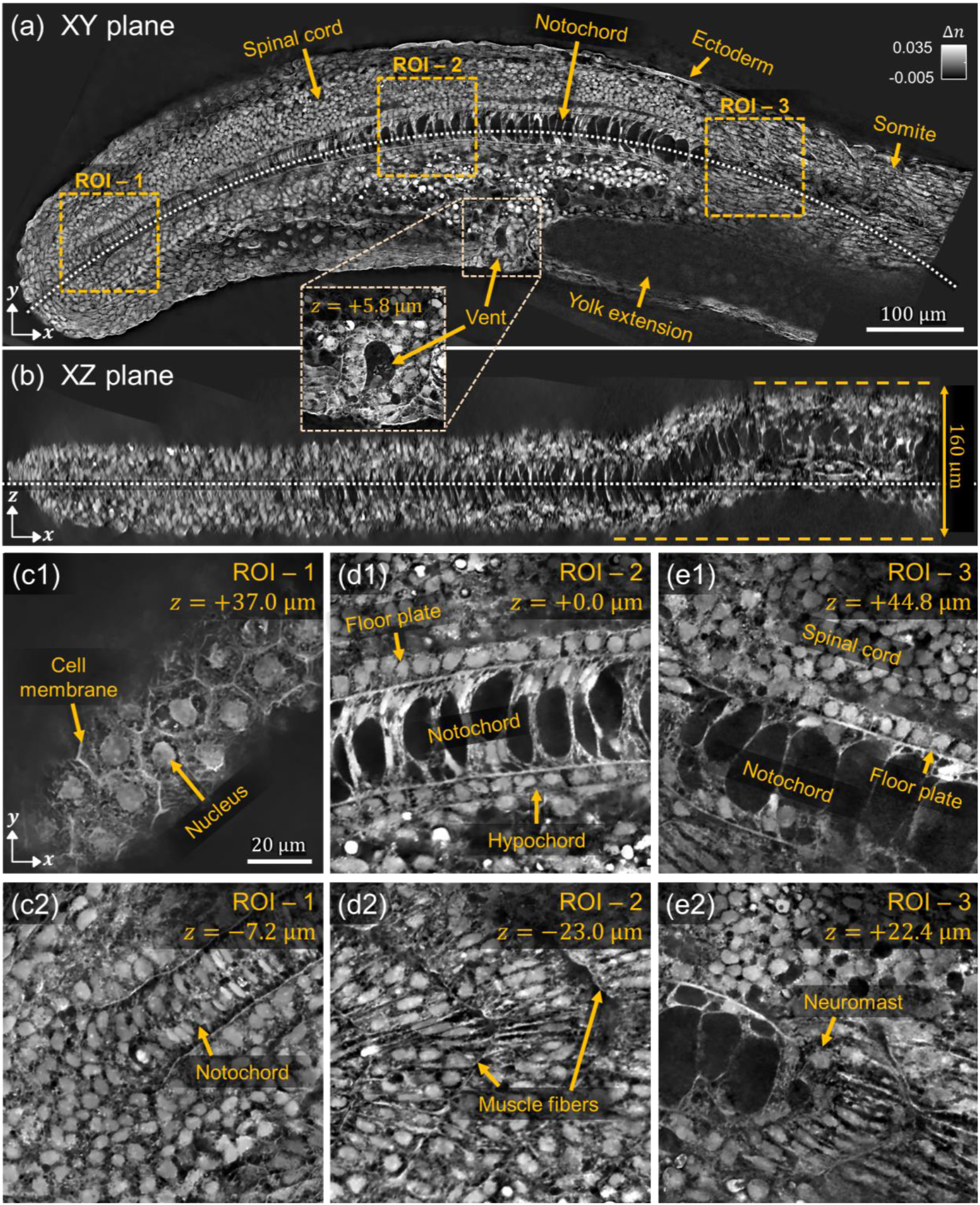
MSBP-based inverse-scattering reconstruction of a 24-hour post-fertilization zebrafish embryo, with large-scale anatomical landmarks highlighted. **(a)** Cross-sectional view of the central XY plane, along with **(b)** a curvature-corrected XZ plane along the dashed white line in (a), reveal the 3D morphology based on RI contrast. **(c1–c3), (d1–d3)**, and **(e1–e3)** show the zoomed-in XY views at different depths within three regions of interest ROI-1, ROI-2, and ROI-3, respectively. These zoomed-in views reveal subcellular-level resolution, enabling clear identification of fine structural features.

Importantly, the region indicated by the arrow in **Figure 8(a)** corresponds to the yolk extension, which could not be successfully reconstructed likely due to its excessive scattering and absorption properties. **Figure S7** in the Supplementary Information shows that the scattering profile of the yolk region consists essentially of chaotic speckle, with speckle patterns from adjacent illumination angles appearing almost completely uncorrelated. Effectively performing inverse-scattering in such highly heterogeneous regions will be an important direction for future research.

## Discussion and conclusion

Our motivation for this work stems from the long-standing fundamental challenge in optical imaging regarding its inability to see deep into tissue due to strong biological scattering. Hardware-based approaches have been developed to physically resist or reverse effects of scattering, but often face challenges such as limited degrees of freedom for scatter-correction, reliance on guidestars, the need for labeling, and high system costs^21-23^. In contrast, computational inverse-scattering methods aim to digitally undo scattering effects. In this approach, the primary challenges lie in the accuracy of the physics-based scattering model and the efficiency of the inversion algorithms. As computational power improves, these challenges become more manageable. However, a fundamental issue remains largely unaddressed: most multiple-scattering inverse solvers are inherently non-convex and can produce different converged solutions depending on the specific implementation strategy, even when using the same underlying physical model to reconstruct from a single dataset. In complex scattering scenarios, these solutions may fail to reflect the sample’s true physiological structure.

In this work, we aimed to develop a methodology for MSBP-based inverse scattering that enabled robust and accurate 3D RI reconstruction in optically thick, scattering samples. Recall that a sample’s 3D RI distribution can itself be interpreted as a volumetric image that enables label-free and quantitative visualization of the sample’s morphology, providing detailed insights into cellular structures, multicellular organization, and tissue architecture. Importantly, RI imaging enables these features to be visualized with high contrast without the need for exogenous labels or perturbative agents. Furthermore, aside from serving as a source of imaging contrast, RI is also the fundamental physical parameter governing a sample’s scattering behavior. Thus, accurately reconstructing RI not only enables morphological imaging but may also have valuable downstream implications for scatter-correction in other techniques, such as wavefront shaping^96,97^, adaptive optics^98,99^, super-resolution^100,101^, and 3D virtual labelling^102-104^. Thus, reconstructing RI in scattering samples may have several important applications for biological imaging.

To formulate an effective inverse-scattering strategy, we first started with 3D scattering microphantoms, which provided ground-truth structures for initial validation and benchmarking. This enabled us to identify implementation strategies that produce robust and accurate RI maps. We then extended and refined these approaches using intestinal organoids, as examples of biologically relevant and more complex scattering samples. Importantly, all implementation strategies were evaluated across consistent datasets, thereby controlling for acquisition conditions, sample content, and signal-to-noise ratio. This allowed for a fair, side-by-side comparison of implementation strategies under consistent conditions.

We found that using amplitude-only measurements in the MSBP inverse solver in Eq. (4) generally produced more stable and biologically realistic 3D RI reconstructions compared to using complex-valued field measurements. This result is surprising, given that field measurements contain both amplitude and phase information, which intuitively should provide more data for reconstruction. We speculate that the phase component introduces ambiguities due to phase-wrapping, which can mislead the non-convex inverse solver and cause it to converge to incorrect solutions. This effect was consistently observed in both phantom and organoid datasets, where reconstructions from field data exhibited strong artifacts that worsened as the level of phase-wrapping increased in the raw field measurements.

Interestingly, the complex field data still indirectly proved valuable. By numerically defocusing the measured fields and then incorporating their amplitude components into the inverse solver, we were able to resolve artifacts that persisted when using only angular amplitude measurements. This suggests that combining angular and defocus diversity in the dataset offers stronger constraints for the inverse solver, helping it converge toward a more accurate and physiologically meaningful solution. Because our system captures full field data, we could generate these defocused measurements computationally. A simpler non-interferometric system could likely be developed that combines angular illumination with high-speed remote focusing to directly capture angular amplitude images at multiple focal planes – thus -achieving comparable reconstruction quality without requiring full complex field measurements.

Several strategies can be explored to further improve the methodology introduced in this work. For example, future work could focus on improving the digital refocusing process, such as determining the most informative focal planes and deciding how many defocus levels to include in order to enhance reconstruction quality. Furthermore, although our datasets in this work composed of scattering measurements resulting from angular plane-wave illumination applied one direction at a time, future research could examine how the choice of illumination angles or the simultaneous use of multiple angles influences reconstruction results^105-107^. Another possibility is to design non-planar illumination wavefronts or structured patterns that effectively constrain the inverse problem and improve robustness^108,109^. In addition, while total variation was used as the regularizer in this work, more advanced forms of regularization could be developed. For example, as realistic bio-mimicking scattering microphantoms become more accessible^110-112^, it may be possible to train neural networks that map scattering measurements to RI distributions for specific types of samples^113-115^. These networks could then serve as learned regularizers that provide more targeted constraints, improving the convergence, and accuracy, and speed of the inverse-scattering process for particular biological systems. Lastly, other studies have also explored using neural networks to significantly accelerate the inverse-scattering process^51,116-118^. Incorporating such approaches could offer valuable improvements in this context as well.

Lastly, all of the inverse-scattering results presented in this work are based on the MSBP model, which is a relatively simple framework for light scattering that assumes forward-only, paraxial propagation. While the practical limitations of this assumption are not yet fully established (e.g., MSBP can provide quantitatively accurate reconstruction even with non-paraxial imaging), it is reasonable to expect challenges as samples become increasingly scattering beyond the regime where MSBP provides accurate modeling. These challenges may include reconstruction artifacts at interfaces with sharp RI transitions (such as with certain organoids as shown in the Supplementary Information, **Figure S7**), as well as failures to reconstruct in regions exhibiting highly chaotic scattering (such as with the yolk region in the zebrafish embryo in **Figure 8**). Scattering models more advanced than MSBP have been proposed and may be better suited for such challenging cases^55,56,119,120^. However, their added complexity makes rigorous evaluation even more critical, as they may exhibit different convergence behaviors and require distinct implementation strategies. Thus, if properly optimized, these models may extend the reach of inverse-scattering methods to a broader class of samples, including to those where our current approach struggles. We hope this work motivates future efforts to systematically refine and adapt such inverse-scattering techniques for deep-tissue imaging in increasingly thick, multicellular samples, paving the way toward robust, large-scale volumetric biological imaging.

## Data and code availability

Data underlying the results presented in this paper may be obtained from the authors upon reasonable request

## Acknowledgements

J. Kim and S. Chowdhury were funded by grants from the National Institute of General Medical Sciences (NIGMS) of the National Institutes of Health (NIH) (R35GM155424) as well as from the Chan Zuckerberg Initiative (2023-321173, 2021-225666). K. Moshksayan, R. Khanna, and A. Ben-Yakar were funded by the National Institute of General Medical Sciences (NIGMS) of the NIH (R01-GM148906) and by the National Institute of Environmental Health Sciences (NIEHS) of the NIH (R43-ES029890). E.S. Cenik was funded by NIH NIGMS (R35GM138340) and Welch Foundation (F-2133-20230405) grants. M.K. was supported by a Welch Foundation grant (F-2008-20220331). M.E. Swartz and J.K.E. were funded by NIH awards R35DE029086, R01AA023426 and R01AA031346. M. Ziemczonok and M. Kujawińska were funded by Warsaw University of Technology, Poland under the program Excellence Initiative: Research University (IDUB). K. Allenspach was supported by internal funding at University of Georgia (UGA).

Wild-type and *daf-2(e1370)* mutant *C. elegans* were provided by the Caenorhabditis Genetics Center, which was funded by NIH Office of Research Infrastructure Programs (P40 OD010440). Consumables for the culture of canine intestinal organoids was supported by internal funding at UGA. We thank Wojciech Krauze and Arkadiusz Kuś for insightful conversations about inverse-scattering.

## Methods

### Optical system

Our experimental system consists of a Mach-Zehnder off-axis interferometer configured to perform angular illumination scans, as illustrated in **Figure 2(a)**. Light from a red laser (Thorlabs CPS635R, 635 nm center wavelength) is delivered via a single-mode fiber into the system after being collimated using lens L1 (Newport MVC-10X Air, NA 0.25). The collimated beam is then split into a sample arm and a reference arm using a beam splitter. In the sample arm, the beam is directed onto a dual-axis galvanometric mirror that steers the beam’s angle. A 4f optical relay composed of lens L2 and a condenser objective lens images the galvo mirror plane onto the sample plane, allowing precise control of the illumination angle. Light scattered from the sample is collected by a second 4f system, consisting of an imaging objective lens and lens L3, and directed toward the detection path. To maintain symmetry and minimize aberrations, we use identical optics in both arms: lenses L2 and L3 are matched (Thorlabs ACT508-200-A), as are the condenser and imaging objectives. In the case when optimizing for high resolution, oil-immersion objective lenses were used (Nikon CFI Plan Apochromat Lambda D 60X Oil, NA 1.42). When optimizing for long working distance to image thicker samples, water-dipping objective lenses were used (Nikon N16XLWD-PF 16X Water, NA 0.8, WD 3.0 mm). Finally, the sample and reference beams are recombined at a second beam splitter, forming a spatially modulated interferogram that is captured by a CMOS camera (ToupTek IUA20000KMA).

### Data acquisition and pre-processing

Interferograms are first acquired at various illumination angles for both the sample and an empty region without the sample (i.e., background). These angles are controlled by steering the 2-axis galvo mirror. From each interferogram, complex-valued field images containing both amplitude and phase are reconstructed by filtering the sample’s spectrum in Fourier space, following standard procedures for quantitative phase imaging via off-axis holography. This yields a set of amplitude and phase measurements across different illumination angles for both the sample and background. Background correction is performed by complex division of the sample and background fields, corresponding to dividing the amplitude and subtracting the phase terms. This step compensates for system-induced phase variations unrelated to the sample. **Figure 2** illustrates this data acquisition pipeline using a scattering microphantom.

Depending on the reconstruction strategy, either the background-corrected complex field measurements or only their amplitude components were used in the MSBP-based inverse-scattering solver described in Eq. (4). The complex field data can also be combined with the Rytov approximation and the Fourier diffraction theorem to analytically reconstruct an initial RI distribution for the sample under the weak-scattering assumption. This initial estimate can serve as a warm start for the MSBP solver to help to stabilize and accurately guide the convergence process, as demonstrated when using field measurements in the inverse solver (see **Figure 4**).

Lastly, complex field measurements can be digitally propagated to synthetically generate defocus diversity without having to physically capture defocused measurements. This was shown earlier to reduce reconstruction artifacts when using amplitude-only data in the MSBP solver (see **Figure 4** and **Figure 5**). To perform digital propagation, a Fourier transform is first applied to the measured field. The resulting spectrum is then multiplied by the free-space transfer function from Eq. (1), followed by an inverse Fourier transform to reconstruct the propagated field.

Technical details on data acquisition and computational reconstruction for the various samples are provided in **Table S1** of the Supplementary Information.

### MSBP inverse-scattering for complex-valued field and amplitude-only measurements

Here, we provide additional details about the mathematics that compose the MSBP inverse-scattering model. As discussed earlier, MSBP models light scattering by representing the 3D sample volume as a series of infinitesimally thin 2D layers, each acting as a phase-modulating element. In Eqs. (2) and (3), we mathematically defined the wave propagation and phase modulation steps that determine how the field at the *m*-th layer, *E*_*m*_, evolved based on the RI at that layer and the field from the preceding layer, *E*_*m*−1_. By starting with the field at the 0-th sample layer (i.e., the illumination field), repeatedly applying Eqs. (2) and (3) propagates the incident field layer-by-layer through the complete sample. The sample’s cumulative scattering effects are ultimately expressed in the output field at the final layer, i.e., *E*_*M*_ for a *M*-layer sample.

In practice, however, *E*_*M*_ does not correspond directly to the field at the imaging plane, which is typically focused to a specific depth within the sample. Furthermore, to physically reach the imaging plane, *E*_*M*_ must propagate through the detection optics and must thus be subject to the finite numerical aperture (NA) of the objective lens, limits spatial resolution. As a result, the field at the imaging plane is a back-propagated and resolution-limited version of *E*_*M*_ and can thus be expressed as:

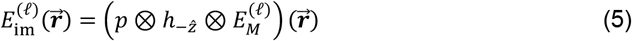

Recall that 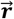 denotes the 2D transverse (*x, y*) spatial coordinates, *ℓ* denotes the index of measurement, and ⊗ represents the convolution operator. The term 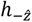 is the optical back-propagation kernel corresponding to the distance 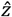 between the final sample layer and the conjugate imaging plane within the sample. Finally, *p* represents the system’s point spread function (PSF), which is determined by the NA of the detection optics and defines the system’s spatial resolution.

If the system directly measures complex-valued fields, the MSBP forward model describing the raw measurement for some known 3D RI distribution is simply:

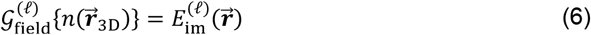

For mathematical clarity, we use slightly different notation here compared to before by introducing the term 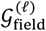 as the MSBP forward model for *complex-valued field measurements*. Recall that 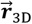 denotes the 3D spatial-coordinate (*x, y, m*), where *m* indexes the axial layers of the sample’s 3D RI distribution, i.e., 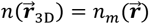 for *m* = 1,2, … *M* for a *M* -layer sample. Note that to measure complex-valued fields, some version of quantitative phase imaging (QPI) must be used. In this work, we computationally extract the complex-valued field term from an off-axis interferogram.

The MSBP forward model for *amplitude-only measurements* can be simply expressed as:

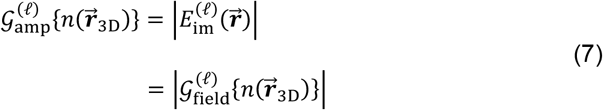

Thus, for a known 3D sample RI distribution, the MSBP forward models in Eqs. (6) and (7) predicts either the complex-valued field or amplitude-only scattering measurements that the imaging system would acquire.

In practice, the goal of inverse-scattering is to infer the sample’s 3D RI from a set of scattering measurements. As described earlier in Eq. (4), we perform this inversion by iteratively minimizing the difference between experimentally measurements and model predictions. For mathematical clarity, we re-express Eq. (4) below:

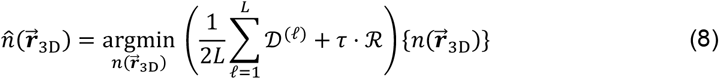

Recall from Eq. (4) that *ℛ* denotes the regularization operator that stabilizes the iterative search. The parameter *τ* sets the strength of regularization and is empirically tuned. Here, we introduce the term *𝒟* to consolidate the least-squares term in Eq. (4) based on whether complex-valued field or amplitude-only measurements are being used:

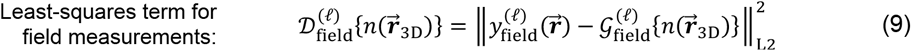

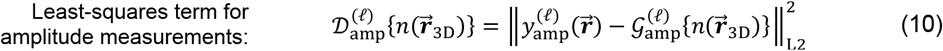

Here, we use the terms 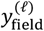 and 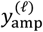 to differentiate between complex-valued field versus amplitude-only measurements.

We solve the minimization problem in Eq. (8) using accelerated stochastic gradient descent. This requires computing the gradient of the least-squares term *𝒟*, either through automatic differentiation or manual derivation. To better understand how the measurement type influences the optimization, we analytically derive the gradient expressions for both the complex field and amplitude-only measurements (see Supplementary Information), as summarized below:

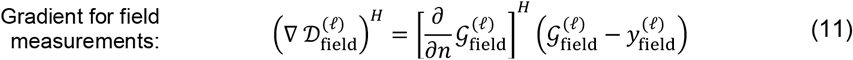

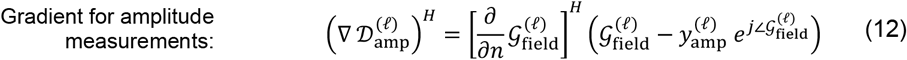

Here, ∇ denotes the gradient operator and *H* represents Hermitian transpose. Interestingly, the gradient expressions above for complex-valued field and amplitude-only measurements are nearly identical and can both be expressed in terms of the MSBP forward model for complex-valued fields. The key distinction between Eqs. (11) and (12) is that the amplitude-only measurement 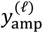 in Eq. (12) is paired with the phase of the predicted field 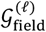. Interestingly, this combined term 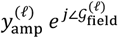 is effectively a synthetic complex-valued measurement that mimics the 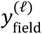 term in Eq. (11). A detailed derivation of these gradients, along with a step-by-step guide for implementing the inverse-scattering framework, is provided in the Supplementary Information.

Notably, while the mathematical difference between the two gradient expressions appears subtle, it leads to substantial differences in convergence behavior and accuracy, as shown in the 3D reconstructions in **Figure 4** and **Figure 5**. Future work could investigate the theoretical implications of this difference and explore whether modified formulations might further improve reconstruction accuracy and stability.

### Quantitative characterization and selection of step size and regularization parameters

To iteratively solve the minimization problem in Eq. (8), users must select values for the step size and the regularization parameter *τ*. We found that these choices significantly influence the reconstructed refractive index (RI) values, impacting the quantitative accuracy of the RI reconstructions. To empirically determine suitable parameters, we collected angular scattering measurements from polystyrene, polymethyl methacrylate, and silica microspheres (BangLabs) of varying diameters, immersed in index-matching oils (Cargille). We then reconstructed the RI maps of the microspheres using a range of step size and regularization values.

Because the true RI contrasts of these samples were known, we were able to evaluate the quantitative accuracy of each reconstruction and identify parameter combinations that yielded the most accurate results. Given the nonconvex nature of the inverse problem, multiple parameter pairs achieved high accuracy on a single sample. To increase robustness, we then selected the parameter combination that consistently produced accurate reconstructions across all tested samples. See **Figure S1** in the Supplementary Information to see more details about this selection process.

Finally, we validated this parameter set using the 40-µm scattering phantom, confirming its generalizability. These parameters were then used for all subsequent reconstructions of biological samples.

### Sample preparation for imaging experiments

#### Microphantom samples

The microphantoms were fabricated via the two-photon polymerization lithography^121^ using the Photonic Professional GT2 system (Nanoscribe GmbH & Co. KG, Karlsruhe, Germany), equipped with 1.4 NA 63x microscope objective, piezo stage for vertical positioning of the sample, and dual-axis galvo for lateral scanning.

The fabrication process leveraged 3D-printing principles, where a tightly focused femtosecond laser was scanned across liquid resin (IP-DIP2, Nanoscribe GmbH & Co. KG, Karlsruhe, Germany). This induced two-photon absorption, polymerizing the resin along the laser’s path. This technique enabled fabrication of microstructures, with high 200 nm lateral resolution, directly on standard #1.5H coverslips. Crucially, the local energy dose could be precisely controlled to dictate the degree of monomer cross-linking, which in turn was proportional to the resulting RI^111,122^. Thus, microstructures were fabricated with carefully calibrated 3D spatial profiles and RI. Moreover, the fabrication process was performed in *dip-in* configuration^123^, where the microphantoms were cured layer-by-layer as the substrate moved away from the objective lens. This strategy prevented the polymerization beam from being degraded by previously cured layers, thereby ensuring the high fidelity of the structures throughout their entire height.

After patterning, unpolymerized resin was removed by a 12-minute bath in PGMEA (Propylene glycol monomethyl ether acetate; Sigma Cat: 484431), followed by a 1-minute bath in N7100 (Novec 7100 Engineered Fluid; Sigma Cat: SHH0002), and then air-drying. Drawing from previous work^74,124^, microphantom designs featured a cell phantom embedded within a pseudo-random distribution of rods. These rods alternated their orientation in successive layers, with each rod measuring 0.5 µm in width and 1.8 µm in height. The lateral spacing between rods in each layer was randomized from 0.7 µm to 3 µm, and layers were stacked vertically every 1.4 µm. The polymer occupied approximately 25% of the total volume (fill-factor), with scattering strength determined by the size of the scattering region (40– 100 µm). During imaging experiments, the microphantoms were immersed in index-matching oil (Cargille, RI = 1.512), which also served as the immersion medium for the oil-immersion objective lens. The objective lens was directly immersed into the same oil, allowing imaging without an additional coverslip on top of the sample.

#### C. elegans worms

Wild-type and *daf-2(e1370)* mutant *C. elegans* were obtained from the Caenorhabditis Genetics Center (CGC). To prepare for imaging experiments, 5-10 wild-type and mutant worms were transferred onto a large coverslip in 50 μL of phosphate-buffered saline (PBS). Excess PBS was carefully removed using a 10 μL pipette tip to avoid disturbing the specimens. The worms were fixed with 50 μL of 4% paraformaldehyde in 1x PBS for 2 minutes, after which the fixative was removed. A graded ethanol rehydration series was then performed: 50 μL of 95%, 75%, 50%, and 25% ethanol were sequentially applied, each for 2 minutes, with the ethanol removed between steps. To complete rehydration, animals were incubated in 50uL of PBS for 2 minutes. After removing the PBS, the animals were resuspended in 40 μL of double-distilled water and remounted on a coverslip.

#### Organoids

Colonic tissue used in this study was obtained post-mortem from a single female research-bred Beagle (∼2 years and 7 months old, 8.4 kg) sourced from Marshall BioResources. The animal was euthanized for reasons unrelated to this study. Tissue collection was performed in accordance with Marshall BioResources’ Institutional Animal Care and Use Committee (IACUC)-approved Standard Operating Procedures, which follow the AVMA Guidelines for the Euthanasia of Animals. All methods, including those for euthanasia and tissue harvest, complied with USDA Animal Welfare Regulations. Culture and maintenance of organoids were performed as per published protocols^125,126^. To prepare samples for imaging, organoids were suspended in Matrigel, seeded as domes into 24-well plates and cultured for 5 days. On Day 5, organoids were recovered from the Matrigel domes by replacing the culture medium in the wells with 0.5 mL of cold Cell Recovery Solution (Corning Inc.). The domes were triturated gently several times, and the contents of the wells were transferred to a 15 mL conical tube. The tube was then filled with 10 mL of ice-cold PBS (1×) and centrifuged at 100 g for 3 minutes at 4°C. The supernatant, containing most of the Matrigel, was aspirated, and the pellet was kept on ice.

For preparing the sample in **Figure *5***, the pellet was resuspended in Matrigel and gently pipetted multiple time to homogenize. Two pieces of stacked tapes, each with a height of ∼450 µm, were attached to either side of a cover slide and a 40 µL drop of the mixture was transferred to the center of that cover slide. The drop was sandwiched by placing another #1.5 cover slide on the top and transferred for imaging

For preparing the sample in **Figure *7***, the organoid pellet was fixed by resuspending it in 1 mL of 4% PFA (Santa Cruz Biotechnology), followed by incubation at 4°C for 1 hour. In the middle of the fixation period, the organoids were gently resuspended once. To validate the thickness of organoids, the nuclei of cells were stained with DAPI (excitation: 402 nm, bandpass: 410–489 nm) after organoids’ recovery from the fixing solution using the same centrifugation setting as mentioned above. A 2× working solution of DAPI was prepared in PBS and added to the organoid pellet, followed by incubation at 4^°^C for 30 minutes. The organoids were washed twice with PBS to remove any unbound DAPI and centrifuged to recover organoids in a pellet. Next, 100 µL of PBS was added to the pellet, pipetted a few times and then the suspension was dispensed at the center of a 50 mm µ-dish (Ibidi, low, polymer coverslip). 4 mL of PBS was slowly added to wet the entire dish area and enable imaging using water-dipping objectives. Finally, the dish containing the fixed and stained organoids was transferred for imaging.

#### Zebrafish embryos

Zebrafish were housed at UT Austin under IACUC protocol AUP-2023-00297. Zebrafish embryos were raised according to [^127^] and staged according to [^128^]. Prior to imaging 24 hours post fertilization (hpf) embryos were fixed in 4% paraformaldehyde overnight at 4 degrees Celsius. Subsequently, they were placed in increasing concentrations of methanol in phosphate buffered saline (PBS) until 100% methanol was reached and embryos were dehydrated. Embryos were then stored at -20 degrees Celsius. To prepare embryos for imaging embryos were rehydrated by placing them in increasing concentrations of PBS in the methanol/PBS solutions until 100% PBS was reached.

#### Spheroids

Cultures of MCF-7 breast adenocarcinoma cells (ATCC, passages 10-15) were maintained in high-glucose DMEM containing fetal bovine serum (10%), Penstrep (1X) and insulin (10 µg/mL) housed in a sterile incubator at 37 °C, 5% CO2 and 99% humidity. For spheroid preparation, cells were harvested using Trypsin EDTA (0.25%) and counted using an automated cell counter (Countess, Thermo Fisher). Cells were then seeded in a Aggrewell 800 plate (STEMCELL Technologies) at a density of 300,000 cells/well. The microwells of the plate were pre-coated with an anti-adherence solution using a standardized protocol, and the plate was centrifuged at 300 rcf for 3 min after cell seeding. MCF-7 spheroids were allowed to form over a period of 12-24 h in the Aggrewell plate in the incubator. For hydrogel embedded samples, such as the one shown in Figure S5, spheroids were collected by gentle agitation using a pipette and centrifugation at 100 rcf for 1 minute. Collagen type IA-based Cellmatrix Cell culturing kit (Nitta Gelatin) was used for embedding the spheroids and gels were cast on a 35 mm Mattek glass bottom dish. On the day of imaging, cells were fixed using 4% paraformaldehyde treatment (gentle rocking for 3 h) followed by washing using PBS. The gels were submerged in PBS for the entire duration of imaging. For suspension samples, such as the one shown in Figure S7, spheroids were collected by gentle agitation using a pipette and centrifugation at 100 rcf for 1 minute. The spheroids were then fixed in suspension using 4% paraformaldehyde by gentle rocking, washed with PBS and dropped onto a glass coverslip for imaging.

